# Automated verification, assembly, and extension of GBM stem cell network model with knowledge from literature and data

**DOI:** 10.1101/2021.07.04.451062

**Authors:** Emilee Holtzapple, Brent Cochran, Natasa Miskov-Zivanov

**Author notes:** University of Pittsburgh, Pittsburgh, PA USA. Tufts University School of Medicine, Boston, MA USA.

## Abstract

Signaling network models are usually assembled from information in literature and expert knowledge or inferred from data. The goal of modeling is to gain mechanistic understanding of key signaling pathways and provide predictions on how perturbations affect large-scale processes such as disease progression. For glioblastoma multiforme (GBM), this task is critical, given the lack of effective treatments and pace of disease progression. Both manual and automated assembly of signaling networks from data or literature have drawbacks. Existing GBM networks, as well as networks assembled using state-of-the-art machine reading, fall short when judged by the quality and quantity of information, as well as certain attributes of the overall network structure. The contributions of this work are two-fold. First, we propose an automated methodology for verification of signaling networks. Next, we discuss automation of network assembly and extension that relies on methods and resources used for network verification, thus, implicitly including verification in these processes. In addition to these methods, we also present, and verify a comprehensive GBM network assembled with a hybrid of manual and automated methods. Finally, we demonstrate that, while an automated network assembly is fast, such networks still lack precision and realistic network topology.

## 1 Introduction

Understanding a disease at a mechanistic level is a complex task, requiring extensive knowledge of how affected genes influence disease progression. Signaling networks are studied to gain more comprehensive understanding of a disease, or to predict potential therapeutic targets [1-3]. The sheer size of the known human interactome [4] drives the need for automated methods for creating and understanding signaling networks. However, current state-of-the-art methods for automation of network assembly are fraught with obstacles that make accurate network assembly time-consuming and labor-intensive [5-7].

Natural language processing (NLP) enables faster information retrieval, but at the price of reduced accuracy. Even manually extracted information may be inconsistent from one source to the next. Biologically accurate interactions are critical for assembling signaling networks, since even one misplaced interaction can have drastic consequences for understanding the true function and behavior of the network.

Current approaches for quantifying signaling network accuracy include validating dynamic network outcomes against experimental data or verifying the individual interactions with experimental data or previous knowledge [8]. For static network models, validating dynamic behavior is not possible. Furthermore, dynamic network models can still match experimental outcomes while relying on interactions that do not happen *in vivo*. These interactions may have been incorrectly inferred from data or existing literature. An informative, effective signaling network consists of individual interactions that accurately depict the system being studied. To verify each individual interaction through experimental work is both tedious and time-consuming, and in many cases, completely redundant since a considerable portion of the human interactome has already been mapped [4]. Many publicly accessible databases collect data on supporting textual and experimental evidence that verify the existence of a putative interaction [9-11]. These databases are programmatically accessible and facilitate the creation of automated tools for network verification.

In this work, we present a methodology for verifying cellular signaling and interaction networks, based on topological features as well as existing data on interactions from several databases. We also show that some of the verification steps can be used in the process of automated model assembly and extension. Using our proposed verification methodology, we evaluate a detailed manually assembled GBM stem cell signaling network, and we compare it with a network that is assembled in a fully automated manner, and with other previously published GBM networks. To this end, the main contributions of this work are:

1. A methodology to verify causal network models of cellular signaling.
2. A verified and comprehensive GBM stem cell network model.
3. An evaluation of automated assembly of context-specific network models from literature.

## 2 Background

While the cell signaling models are typically used to study the dynamics, and they rely on either ODEs [12], reaction rules [13], or element update rules [14-16], in this work, we focus mainly on the underlying structure of these models, a graph *G*(*V, E*) with a set of nodes *V* and a set of edges *E*. The graph nodes represent biological system components such as genes, proteins, small molecules, miRNAs, and biological processes. We refer to these biomolecules and processes as “entities”. The graph edges represent physical or functional relationships between entities, which we refer to as “interactions”. Interactions with a known cause-effect relationship form a directed graph, while the interactions where the nodes are known to have a relationship, but the cause and effect is unknown, form an undirected graph. Depending on the available knowledge, modeled interactions between elements can be direct, commonly with known mechanisms of interaction (e.g., phosphorylation, binding, methylation), or indirect (causal only), when we know that there is influence, but we do not have detailed knowledge about mechanisms of interactions.

To create signaling network models, a manual assembly process outlined in Figure 1 is commonly used. The first step is a selection of relevant biological system components, that have been shown to play a role in the disease of interest. These components are supplied by a number of sources, including expert knowledge or different publicly accessible databases. These databases may curate canonical signaling pathways, such as KEGG [17] or PANTHER [18], or they may collect data on individual interactions, such as STRING [9] or BioGRID [19]. Interaction databases provide curated, often high-confidence data on signaling pathways in disease or normal cell conditions [20]. For a disease network, we can supplement our network with experimental data that is cell- or patient-specific, such as genes or proteins that have differential expression, somatic mutations, or altered signaling capacity. The final list of biological molecules composes the nodes of the network. The edges are created between nodes based on existing data from literature and interaction databases.

**Figure 1.**
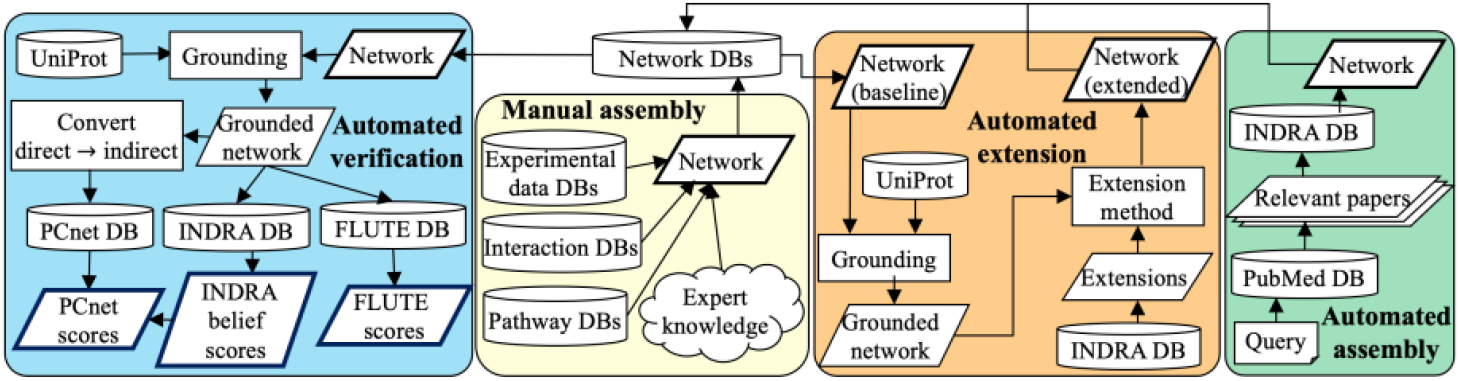
Workflow of the three automated approaches presented in this work, automated network verification, automated network extension, and automated network assembly, and the key sources for manual assembly approach. We illustrate sources of information (data, literature, pathways, interactions, networks) in the form of database (DB) blocks.

## 3 Methodology

In the following section, we describe our methodology for network verification, assembly, and extension.

### 3.1 Automated network verification

Our network verification method, outlined in Figure 1, utilizes the information from several databases to confirm the network structure, by retrieving information about network nodes, retrieving relevant literature, and providing support for interactions.

As shown in Figure 1, the main input for the network verification method is a model network, that is, a list of all its entities and interactions. To use the databases, we first ground entities by finding their unique identifiers (IDs), and this process is dependent on entity type. For genes and proteins, our network verification method utilizes the UniProt API [21, 22] to automatically determine their standard IDs. Specifically, we ground gene and protein names by mapping to both the official gene symbol (OGS) and the Human Genome Nomenclature Committee (HGNC) identifier. These two ID types are very common and allow for comparison to many online resources. The automated ID mapping for genes and proteins allows for fully automated verification of gene regulatory networks, protein-protein interaction networks, and in general, many cell-signaling networks. However, there is no API that would allow fully automated grounding of small molecules, biological processes, and miRNAs. Thus, finding standard IDs for these entities is done manually through database web interfaces such as miRbase [23], Chemical Entities of Biological Interest (CHEBI) [24], or Gene Ontology (GO) [25, 26]. Since proteins and genes tend to be the most common and well-studied endogenous biomolecules, most networks in model databases have fewer chemical, biological process, or mRNA entities. For example, STRING [9], one of the largest interaction databases, contains only genes and proteins. Likewise, BioGRID [19] contains over 2 million protein-protein or protein-gene interactions, while containing <30,000 chemical interactions. If an entity is not matched to a standardized ID from these databases, the plain text name is used instead.

Once all network nodes are grounded, we use three online resources, the INDRA database [5], PCnet [20], and FLUTE [27], to verify the network edges. INDRA is a system that draws on natural language processing tools and structured databases to collect statements about mechanistic and causal entity interactions. INDRA relies on a number of machine readers (e.g., TRIPS [28], REACH [29], etc.) to extract these interactions from literature, and provides a belief score for each interaction in its database. The INDRA statements can include direction information where one entity influences the other (directed), or only indicate that entities are known to interact, but the direction is unknown (undirected). The statements may also include the sign of the influence on the downstream entity. The positive sign corresponds to an increase in activity or amount of the downstream entity, and the negative sign corresponds to a decrease in its activity or amount. The INDRA database also contains statements from papers that have already been processed, which reduces the amount of time required for extracting interactions from large text corpuses. In addition to statements obtained from machine readers, INDRA also provides statements extracted from interaction databases. Finally, the sheer size of the INDRA database (over 2 million statements), as well as its daily updates, make it a useful resource for model verification.

Our network verification method automatically compares grounded model networks to INDRA statements. This step takes one input parameter, the type of network (directed or undirected). For directed networks, it iterates through all interactions in the network, and retrieves all statements that match the entity identifiers, as well as the direction and sign. For undirected networks, it retrieves statements that match the entity identifiers without checking direction or sign. The output of this step are INDRA statements that support interactions in the input network. We also assess the confidence of all interactions in the network using the INDRA belief score [5]. This score, calculated by INDRA, is based on the prevalence of similar statements in literature or observed databases. The belief score ranges from 0.0 (no supporting evidence) to 1.0 (large amount of supporting evidence).

The Parsimonious Composite Network (PCnet) [20] is a high-confidence network of protein-protein interactions. PCnet uses 21 different human interaction databases to inform the network, where each interaction must be found in at least two of the 21 networks. This composite network excludes interactions that are not reproducible, and therefore, it contains only high-confidence interactions. By comparing model interactions to PCnet interactions, we identify which interactions in the model have support from multiple curated signaling networks. While PCnet interactions are highly supported, they are undirected, and independent of context. This distinguishes PCnet from INDRA, which contains many interactions that are directed and contextual, but have lower confidence.

Besides automatically comparing the input network with INDRA, our verification method also automatically compares the network to all interactions in PCnet. Here, we use a local copy of PCnet, stored as a plain text file, which is also freely accessible and available for download from nDex [11].The first step in this case is a conversion of the directed network to an undirected one, since PCnet contains only undirected interactions. Our method then iterates through all model interactions and compares them to all PCnet interactions. Finally, it outputs a list of all interactions within the signaling network that are verified by PCnet. We can verify a network using only PCnet, or we can use PCnet in addition to INDRA.

The Filter for Understanding True Events (FLUTE) [27] tool utilizes existing interaction database resources to find support for machine-read interactions. The FLUTE database stores information from five interaction databases, including protein-protein interactions (PPIs), protein-chemical interactions (PCIs), and protein biological process interactions (PBPIs). For PPIs and PCIs, FLUTE utilizes the STRING interaction score [9], STITCH interaction score [30], or the GO Term annotation [25, 26], to assess the confidence of the interaction, depending on the interaction type. The STRING and STITCH interaction scores are composed of several evidence types to support an interaction, such as experimental data, text mining, or database co-mentions. FLUTE is capable of selecting high confidence interactions and filtering out many incorrectly read interactions within a reading set. FLUTE encompasses several types of biological entities including proteins and genes, chemicals, and biological processes, while PCnet is composed of only proteins. FLUTE is also able to provide a score for the interaction confidence, unlike PCnet. In contrast to INDRA, the FLUTE database contains interactions with a high level of human oversight. FLUTE provides an extra level of scrutiny over INDRA, while still being less restrictive than PCnet. The FLUTE tool was specifically designed for interaction filtering, unlike PCnet, which is a network. In Table 1, we summarize the characteristics of the three sources, INDRA, PCnet, and FLUTE, that are used in our automated verification method.

**Table 1.**
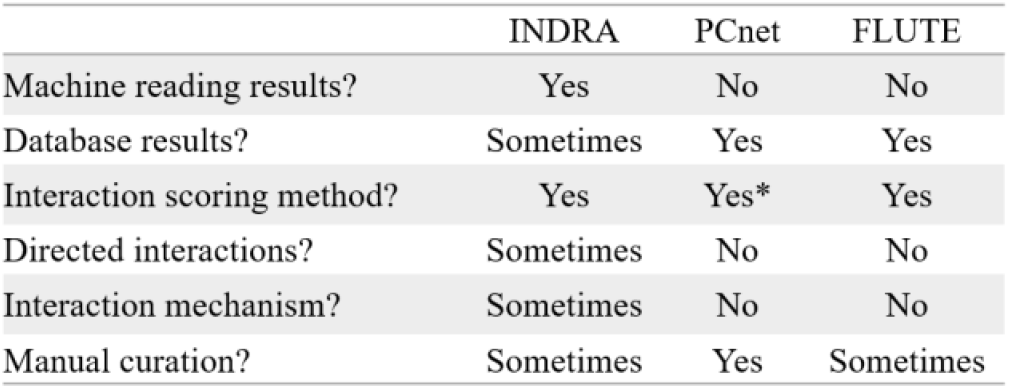
A comparison of the characteristics of INDRA, PCnet, and FLUTE. *The presence or absence of an interaction in PCnet can be used as a binary score.

### 3.2 Automated network assembly

Our automated network assembly method has two main steps, literature search in PubMed [31], followed by the use of INDRA API to retrieve all statements. To assemble a network that includes relevant, commonly affected pathways for a given context (e.g., disease, cell type, tissue, or a biological state), we use a query as an input to the network assembly method. This query has two parts connected with logical AND, and each part is a list of terms connected by logical OR operator. One list of terms leads to the retrieval of papers from the desired context (e.g., “‘Type 1 diabetes’ OR ‘juvenile diabetes’ OR ‘insulin-dependent diabetes’”), and the other list of terms refers to relevant signaling networks and pathways (e.g., “signaling OR network OR pathway OR cascade OR interaction OR regulation”). We note here that other logical expressions are possible [32], and could be supported by our proposed workflow; however, this is beyond the scope of the work presented here. For well-studied contexts or broad queries that return a large number of papers (in the order of 10,000+), we select the most relevant papers, as determined by the PubMed’s Best Match feature [33]. The second step of our automated assembly method takes as input the standardized paper identifiers for the set of context-specific papers found in the first step and searches the INDRA database using its API to find all statements (i.e., interactions) in this paper set.

We limit the returned statements to those that have only two distinct entities. This eliminates statements that represent an edge joining more than two nodes, such as a statement describing complex formation. We restrict the number of entities in a statement in order to be able to compare all statements to PCnet and other existing networks, which represent steps in a signaling pathway as one-to-one interactions. As a result, the automatically assembled network is composed entirely of INDRA statements, where the entities are the nodes, and the interactions stored in the INDRA statement are edges.

To improve the quality of the assembled network, we added several filters to the automated assembly method, including those that we used in our network verification approach. These filters are based on the information included in each INDRA statement. First, the belief score can be applied as a cut-off, since it is determined with respect to the amount of evidence to support the interaction; an interaction with a higher belief score is less likely to be a false positive (an invalid or non-existent relationship between two entities). To filter by belief score, we used the cut-off for a high confidence interaction as described in [5], which is 0.85, and discarded all interactions below that score. Next, we can also filter by interaction type, that is, direct or indirect, as stated by INDRA. For some interactions, INDRA contains evidence on whether an interaction is direct or indirect. Other statements do not contain any evidence on the interaction type. By selecting for only direct interactions or those with high belief scores, the final network contains fewer low-confidence interactions.

### 3.3 Automated network extension

The methods used for automated network verification can also be utilized in the process of network extension, thus providing verified interactions by construction. In cases where an existing (baseline) model, i.e., its underlying network, fails to capture the full detail of the studied system, an automated extension method retrieves new interactions (extensions) to improve network scope. Our goal is to explore whether our proposed methods can help identify these new important entities and interactions to be included in the baseline network.

To select potential network extensions, we again utilize the INDRA DB. First, we take the list of all baseline network nodes, with standardized IDs, and call the INDRA API for each node. When only one entity name is used in the API call, this retrieves all INDRA statements where the entity is mentioned. This gives us a list of extensions, which can either add new nodes to the network or provide edges between unconnected nodes in the baseline network.

Here, we use two different extension approaches. First, we use a simple approach that adds to the baseline network those interactions output by INDRA with at least one entity already in the baseline network. Thus, we know that these extensions will all connect to the network. We also use the TopX extension method, where X refers to the number of extensions added to the network. In general, the extensions can be ranked with respect to any metric of interest. In this work, we will be adding extensions based on their INDRA belief score. The TopX method has two benefits over simple extension. First, the number of extensions added can be tailored to the size of the network. This ensures that the characteristics of the original network are kept and not overshadowed by the sheer number of extensions added. Secondly, when interaction confidence or a belief score is the criteria used for interaction ranking, the TopX method selects and adds only high confidence interactions.

## 4 Results

Here, we first describe a manually created and carefully curated network of GBM stem cell signaling. We will verify the interactions in this network with our proposed automated verification approach. Furthermore, we will compare this manually created network with GBM networks that are automatically assembled using existing databases. We will also use our automated network verification methodology to compare this network with other previously published networks. Finally, we will discuss further extensions of this network that are found using our automated network extension approach.

### 4.1 GBM stem cell network

We assembled the GBM^expert^ network manually, using several different sources. We started building a small initial network by including the key GBM pathways identified from the TCGA mutation data, as well as the key pathways that were found to play important roles in apoptosis, cell proliferation, and DNA damage, in particular, the Rb/CDK [34], PI3K/Akt [35], JAK/STAT [36], Raf-MEK-ERK [37], EGFR [38], p53 [39, 40] Hedgehog [41], and Notch [2] pathways. To encompass a wider selection of cancer-associated pathways and individual elements (oncogenes, tumor suppressors, etc.), we obtained more information from the GBM stem-cell specific literature and added new entities and interactions to the small initial network. The GBM^expert^ network is generic with respect to different GBM stem cell lines, and it includes 134 elements, and 279 interactions between these elements (Supplemental Table 1). Finally, from the GBM^expert^ network, we also created the GBM^expert^_PCnet_ network, which contains only interactions that are supported by PCnet. The sizes of both networks are listed in Table 2.

**Table 2.** Statistics for all studied networks. The clustering coefficient is the average clustering coefficient for all nodes in the network. We calculate the number of hub nodes as those with >7 edges, as well as the relative frequency of hub nodes in each network.

| Network | Node # | Edge # | Average Clustering Coefficient | Connected Components | Hub Nodes |  |
| --- | --- | --- | --- | --- | --- | --- |
|  |  |  |  |  | # | % |
| GBM<sup>expert</sup> | 134 | 279 | 0.07 | 1 | 18 | 13.43 |
| GBM<sup>expert</sup><sub>PCnet</sub> | 118 | 207 | 0.06 | 1 | 11 | 9.09 |
| GBM<sup>query</sup> | 831 | 2388 | 0.08 | 20 | 134 | 16.13 |
| GBM<sup>query</sup><sub>PCnet</sub> | 250 | 412 | 0.05 | 19 | 23 | 9.16 |
| GBM<sup>query</sup><sub>d</sub> | 10 | 6 | 0.00 | 4 | 0 | 0.00 |
| GBM<sup>query</sup><sub>d,u</sub> | 194 | 180 | 0.00 | 37 | 6 | 3.09 |
| GBM<sup>query</sup><sub>0.85</sub> | 269 | 454 | 0.05 | 14 | 20 | 7.44 |
| JeanQuartier20 | 538 | 911 | 0.24 | 5 | 26 | 4.83 |
| TCGA<sup>miRNA</sup> | 278 | 2287 | 0.00 | 1 | 207 | 74.46 |
| GBM<sup>SIGNOR</sup> | 26 | 46 | 0.10 | 1 | 11 | 42.31 |
| Tunc16 | 191 | 242 | 0.043 | 1 | 8 | 4.29 |
| GBM<sup>KEGG</sup> | 45 | 47 | 0.10 | 3 | 0 | 0.00 |

### 4.2 Verification of GBM^expert^

Using our automated network verification method described in Section 3.1, we verified the GBM^expert^ network. In Figure 2a, we show the overlap between the INDRA DB, GBM^expert^ network, and PCnet, the intersection between each two of them, as well as the intersection between all three (GBM^expert^_PCnet_). Each of the 279 interactions in the GBM^expert^ network is found in INDRA, confirming the existence of these interactions, as well as their direction and sign. Consequently, all model interactions in GBM^expert^ that are supported by PCnet are also present in INDRA, and they form GBM^expert^_Pcnet_. Thus, the set of 208 interactions in GBM^expert^_PCnet_ confirmed by PCnet and INDRA, indicates that the majority of interactions in GBM^expert^ are both high-confidence and have a supported mechanism.

**Figure 2.**
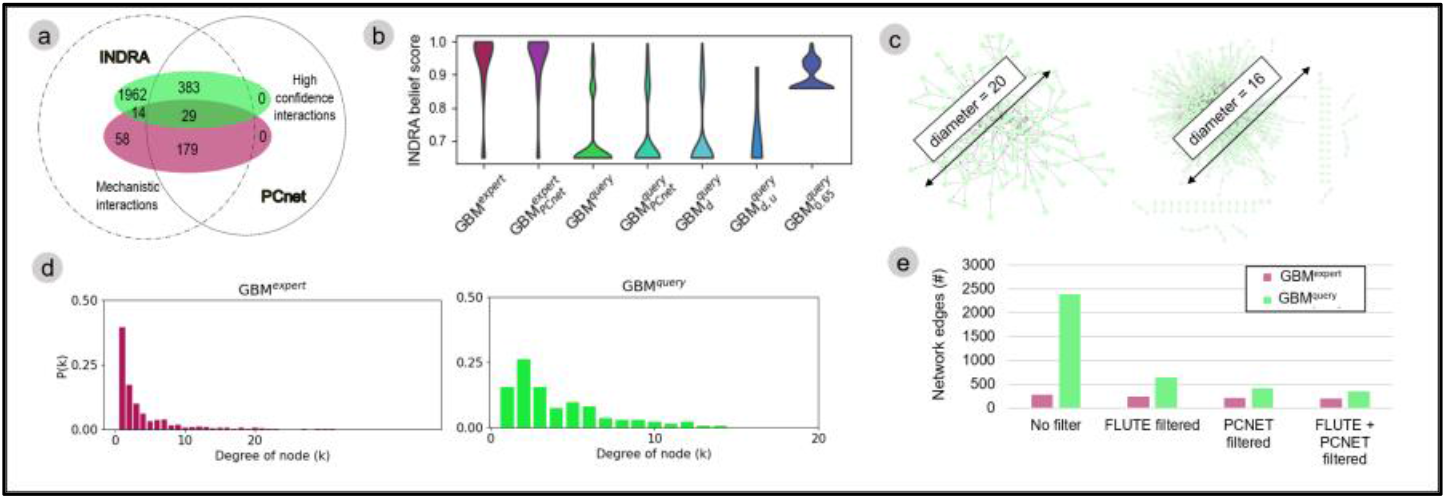
Edge overlap between PCnet, INDRA, GBM^expert^ (maroon), and GBM^query^ (green) (a), the distribution of INDRA belief scores for all manual and automated networks (b), the network diameter for GBM^expert^ (left), and GBM^query^ (right) (c), node degree distribution for GBM^expert^ (maroon), and GBM^query^ (green) (d), and the effect of different filters on network size (e).

As described in Section 3.1, besides finding the existence of the GBM^expert^ network interactions in literature and databases, our verification method also measures the amount of supporting information for individual interactions in the GBM^expert^ network using the INDRA belief score. We show the distribution of INDRA belief scores for each interaction in GBM^expert^, as well as for interactions in GBM^expert^_PCnet_ in Figure 2b. It is important to note that the minimum belief score, if the corresponding statement can be found in the INDRA database, is 0.65. While an interaction that is not in the INDRA database would be assigned a belief score of 0.00, all interactions in GBM^expert^ are found in INDRA. Intuitively, when we remove interactions that are not found in PCnet, the average INDRA belief score increases. This can be attributed to the fact that interactions in PCnet have at least two independent networks supporting each interaction.

We also explore properties of the network, independent of node and edge identity. While these numbers alone cannot verify our network, they can provide us with a measure of how useful the network is, and whether it resembles a signaling network. In Figure 2c (left), we show the GBM^expert^ network figure, as well as the network diameter. GBM^expert^ is one connected component (Table 2), with a network diameter of 20. These numbers confirm that our preliminary baseline network is well connected, without any disconnected nodes or isolated clusters. The network nodes form long paths, typical for signaling networks, instead of star-like clusters.

We also list in Table 2 the average clustering coefficient, and the number and frequency of hub nodes for the GBM^expert^ and the GBM^expert^_PCnet_ networks. The clustering coefficient is a metric of the connectedness of each node within a network [42]. The clustering coefficient for both networks is more indicative of a network that describes a real-world phenomenon than a randomly generated one [43]. Cancer signaling networks depend on the existence of hub nodes, which are highly susceptible to chemical inhibition. We define a hub as a node with >7 edges, which includes both incoming and outgoing edges. Both manually assembled networks show a hub node frequency of approximately 1 in 10 (9-13%).

We also evaluate the node degree distribution in

Figure 2d. Biological signaling networks are thought to be weakly scale-free, where the node degree distribution conforms to a power law. Most nodes will have few interactions, but there are also hub nodes with many in- or out-going edges. We find that our manual baseline network shows scale-free properties. Overall, our verification method and graph-based analysis highlight that our manually created context-specific network is not only composed of high-confidence interactions, but it also conforms to known graph topological characteristics of biological signaling networks.

### 4.3 Automated network assembly

In contrast to the manually created and curated GBM^expert^ network, our fully automated assembly method (described in Section 3.2) created another context-specific GBM network, GBM^query^. To compile a relevant literature set, we performed a targeted search of GBM literature in PubMed, and retrieved the top 600 most relevant papers (according to the PubMed Best Match algorithm [33]), using the following query:

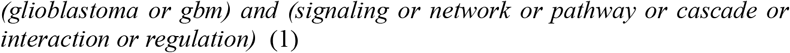

We chose the second term to limit the frequency of clinical or morphological GBM studies, and to retrieve only papers on cell pathways and signaling in GBM. Our automated method obtained all associated INDRA statements from these papers, as described in Section 3.2, and automatically assembled the GBM^query^ network.

To explore whether additional automated filtering methods could help select a useful and reliable GBM network, we analyzed not only the baseline GBM^query^ network, but also its subnetworks. In particular, we explored several subnetworks of the GBM^query^ network that contain more supported or higher confidence interactions. We automatically created these four subnetworks of the GBM^query^ network using three interaction metrics - interaction type (direct or indirect), INDRA belief score, and PCnet overlap (Table 2). It is important to note that unlike GBM^query^, we did not create GBM^expert^ subnetworks filtered by belief score or interaction type due to the high average belief score, as well as the absence of indirect or nonspecific interactions.

We used the interaction type attribute included in INDRA statements, to select only direct interactions within GBM^query^, forming GBM^query^_d_. The reason behind selecting direct interactions is that usually there exists more evidence about them and their mechanisms in literature. However, due to the scarcity of direct interactions in machine reading output, we also created a subnetwork GBM^query^_d,u_, which includes both direct and “unspecified” (when interaction type is not listed in INDRA) interactions from GBM^query^. We created two more networks from the GBM^query^ network by either selecting only those interactions that are present in PCnet to create the GBM^query^_PCnet_ network, or by selecting those interactions with INDRA belief score larger than 0.85 to create GBM^query^_0.85_. We list in Table 2 the size, clustering coefficient, the number of connected components, and the number of hub nodes for the GBM^query^ network and its four subnetworks.

### 4.4 Evaluation of automatically generated networks

Although the five automatically generated networks were assembled using the information from INDRA, the INDRA belief scores of the interactions in those networks can vary significantly (Figure 2b). Compared to GBM^expert^, GBM^query^ has much lower average belief score. Even by restricting interactions from GBM^query^ to belief score >0.85, which excludes interactions found in a single paper, their average belief score is still lower than for GBM^expert^ (0.89 vs. 0.94).

Furthermore, when compared to its subnetwork GBM^query^_PCnet_ (Figure 2a), we found that out of 2388 interactions in the GBM^query^ network, only 412 are represented in PCnet. In other words, PCnet considers over 75% of the GBM^query^ interactions as low-confidence interactions. Furthermore, we find that the network is composed of 20 connected components (Table 2). This suggests that the automated network assembly may not be optimal in terms of interaction quality. However, we analyzed the scale-freeness of GBM^query^ by its node degree distribution (Figure 2d), and the node degree distribution strongly conforms to a scale-free distribution, indicating that GBM^query^ has a network structure typically seen in biological networks. We find that the node degree distribution more closely resembles a power law distribution than a normal distribution, which would be expected for a randomly generated network [44]. We see that the most common node degree in GBM^query^ is *k*=1, which is a degree of input and output nodes.

We also compared the size of manually and automatically created networks. As expected, GBM^expert^ is much smaller than GBM^query^. Due to the importance of hub nodes in cancer signaling, we identified hub nodes in all networks (node degree >7). While GBM^expert^ has less hub nodes than GBM^query^, the proportion of hub nodes is comparable given the size of both networks (13.41% and 16.13%, respectively). Any filtering, for either manually or automatically created network, results in a decrease in the proportion of hub nodes in the filtered network. For instance, selecting only interactions supported by PCnet decreases hub node proportion to 5.31% in GBM^expert^_PCnet_ and 5.58% in GBM^query^_PCnet_. Compounding the effort to improve the networks is the increase in number of disconnected components after filtering. By adding a filter to increase interaction support, we increase the disconnect between network modules. For example, after filtering indirect interactions out of GBM^query^, we are left with 37 distinct, unconnected modules.

Finally, we compare the effect of different verification techniques on the number of edges of GBM^query^ and GBM^expert^ (Figure 2e). GBM^query^, which is much larger than GBM^expert^, is the most affected by filtering. Using either PCnet or FLUTE to filter GBM^query^ reduces the size of the network by more than 1500 edges. However, for GBM^expert^, filtering eliminates at most 75 interactions. FLUTE is the least stringent filter for both networks, while PCnet filters out the most interactions from both networks.

### 4.5 Existing GBM network verification

We compared the literature support of the GBM^expert^ network versus other published GBM networks. We examined two networks publicly accessible from literature, JeanQuart20 (Jean-Quartier et al. [45]) Tunc16 (Tuncbag et al [46]), and two networks from databases GBM^KEGG^ (KEGG[17]) and GBM^SIGNOR^ (SIGNOR [47]). Additionally, we retrieved from nDex [11] a TCGA RNA-miRNA interaction network, TCGA^miRNA^. It should be noted that TCGA^miRNA^ is not a mechanistic signaling network like the others; rather, it is a correlation network derived from gene expression data. Similar to other networks, we summarized in Table 2 the characteristics of these five existing GBM disease networks.

In Figure 3a-d, we show the INDRA, PCnet, and GBM^expert^ overlap with JeanQuart20, Tunc16, GBM^KEGG^ and GBM^SIGNOR^, respectively. We find that, while GBM^expert^ outperforms existing GBM networks in terms of INDRA representation, PCnet representation is more comparable. JeanQuart20 has the highest percentage of interactions represented in PCnet. TCGA^miRNA^ is again, the least supported, since it is a correlation network of gene-miRNA interactions, which has no presence in PCnet, and only infrequent mentions in INDRA.

**Figure 3.**
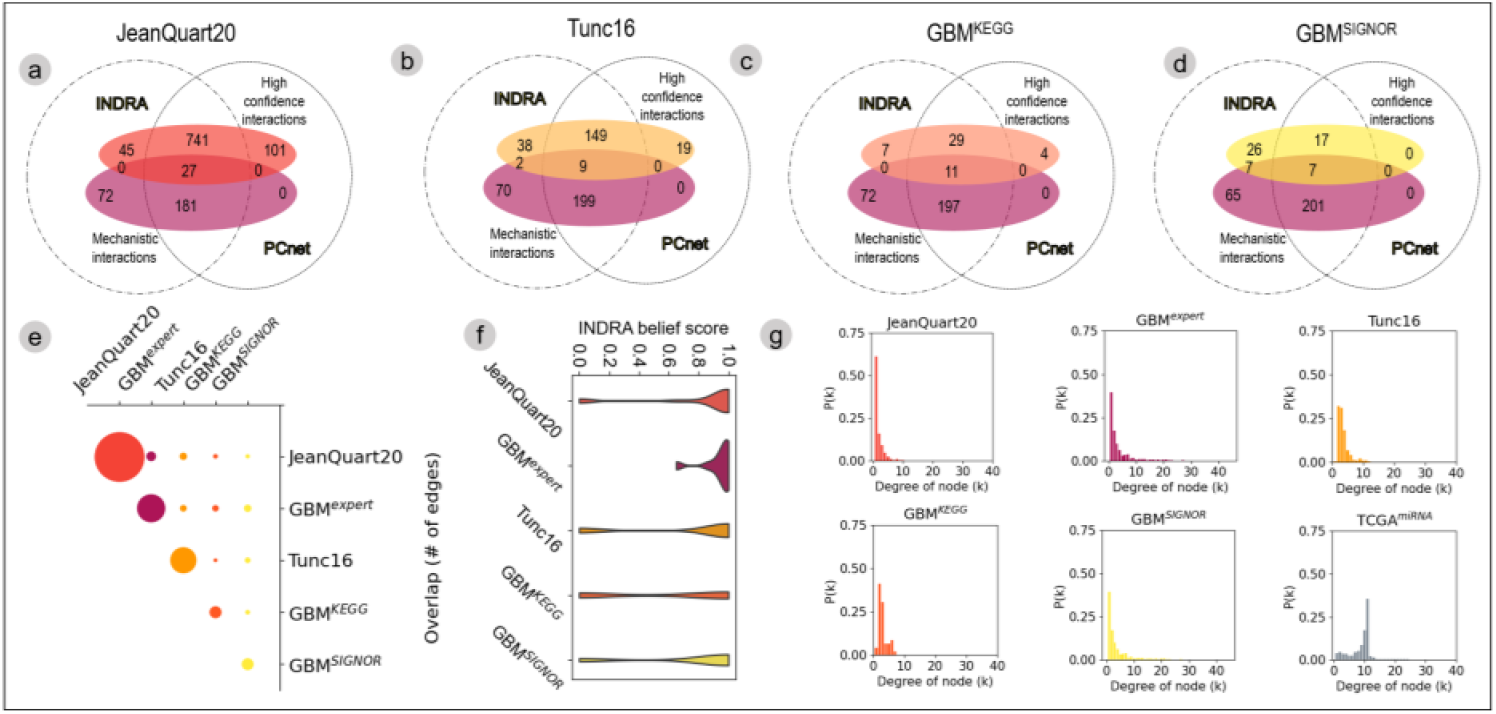
INDRA, PCnet, and GBM^expert^ overlap with JeanQuart20 (a), Tunc16 (b), GBM^KEGG^ (c) and GBM^SIGNOR^ (d). The edge overlap between manually created networks (e), INDRA belief score distribution (f), and node degree distribution for GBM networks (g).

Additionally, we compared the overlap between each pair of GBM networks in terms of shared interactions (Figure 3e). While all five networks are intended to address the same disease signaling network, we find that there is very small overlap. For example, the maximum overlap is between GBM^expert^ and JeanQuart20, and even this overlap is only 28 interactions, making it 10.04% of the GBM^expert^ network and 3.07% of the JeanQuart20 network. This disparity is most likely due to differences in represented pathways.

Using our automated network verification approach (Section 3.1), we evaluated individual interactions within networks (Figure 3f). Again, we find that our proposed GBM network, GBM^expert^, has much higher literature support than other networks, even compared to other manually assembled networks. GBM^expert^ is the only manually created network with all statements supported by the INDRA database, and has a higher average belief score than any other studied network. This is independent of the curation source (i.e., literature or database). These results indicate that the GBM^expert^ network may be less likely to contain interactions that are false positives.

Similar to the networks discussed in previous sections, we calculated the node degree distribution for each of the published networks (Figure 3g). We find that all networks display some degree of scale-freeness. However, TCGA^miRNA^ does not – the most common node degree is 10, which does not conform to the expected power law distribution. This can be explained by the fact that this a correlation network, and so each node is at least weakly connected to many other nodes, which is common for correlation networks [42].

### 4.6 Automated extension of GBM^expert^

Using the methods described in Section 3.3, we retrieved 116,535 edges that can be used to extend GBM^expert^. Using the simple extension method, the manually created baseline model was extended to include approximately 8,000 nodes, which is approximately a 60-fold increase compared to the original network. While this extended network includes many new interactions, it overshadows the original GBM^expert^ network, and is likely populated with interactions and pathways not relevant to GBM. By construction, this extended network includes interactions that are connected with the original GBM^expert^ network, and therefore, it does not suffer from the issue that the GBM^query^ network has – this extended network has only one connected component and a high average INDRA belief score (0.91). While the extensions have a high average belief score, not all are relevant to GBM. For a GBM model that is both high-confidence and GBM-specific, manual selection of retrieved extensions can improve model relevancy.

We also obtained results for extension of GBM^expert^ using the TopX extension method. For X=25, the new interactions prior to model extension were hanging nodes, i.e., an edge between one node in the network and one or two new nodes. For X=50, the extensions are grouped into star-like clusters (4-6 nodes), where a single new node connects to several nodes already in the network. Thus, using automated methods for network extension, we can create high confidence networks that retain structural characteristics of typical biological networks.

## 5 Conclusion

Verification of disease network models is an important component of modeling in systems biology. Publicly available databases are a valuable resource for both model creation and verification, and in this work we focus on automating the use of these databases. We propose a workflow that relies on publicly available resources and includes steps for automated network verification, as well as methods that utilize those steps and resources to automate network assembly and extension. As such, our workflow not only enables more efficient creation of high confidence disease models, but also contributes to systematic reuse and reproducibility of existing information. Additionally, we present and verify a manually created network model of GBM stem cells and compare it with an automatically assembled network. We find that, while both manually and automatically created networks replicate some characteristics of biological networks (i.e., scale-freeness, hub node frequency, etc.), the manually created network has higher-confidence interactions. Using FLUTE, INDRA, and PCnet to verify network interactions, we show that our manually created GBM^expert^ network is composed of well-known, mechanistic interactions. Our analysis holds up, even when comparing our manually created GBM network to other previously published GBM networks. We find that the interactions within our network are supported by more literature and database sources, on average, than the other existing networks.

## Supporting information

Supplemental Table 1 (Expert-curated GBM model logic)

## Acknowledgments

Funded by Defense Advanced Research Projects Agency (W911NF-17-1-0135). Thanks to Yasmine Ahmed and Casey Hansen for helpful discussions.

